# Continuous Tracking using Deep Learning-based Decoding for Non-invasive Brain-Computer Interface

**DOI:** 10.1101/2023.10.12.562084

**Authors:** Dylan Forenzo, Hao Zhu, Jenn Shanahan, Jaehyun Lim, Bin He

## Abstract

Brain-computer interfaces (BCI) using electroencephalography (EEG) provide a non-invasive method for users to interact with external devices without the need for muscle activation. While noninvasive BCIs have the potential to improve the quality of lives of healthy and motor impaired individuals, they currently have limited applications due to inconsistent performance and low degrees of freedom. In this study, we use deep learning (DL)-based decoders for online Continuous Pursuit (CP), a complex BCI task requiring the user to track an object in two-dimensional space. We developed a labeling system to use CP data for supervised learning, trained DL-based decoders based on two architectures, including a newly proposed adaptation of the PointNet architecture, and evaluated the performance over several online sessions. We rigorously evaluated the DL-based decoders in a total of 28 human participants, and found that the DL-based models improved throughout the sessions as more training data became available and significantly outperformed a traditional BCI decoder by the last session. We also performed additional experiments to test an implementation of transfer learning by pre-training models on data from other subjects, and mid-session training to reduce inter-session variability. The results from these experiments showed that pre-training did not significantly improve performance, but updating the models mid-session may have some benefit. Overall, these findings support the use of DL-based decoders for improving BCI performance in complex tasks like CP, which can expand the potential applications of BCI devices and help improve the quality of lives of healthy and motor-impaired individuals.

**Significance Statement:** Brain-computer Interfaces (BCI) have the potential to replace or restore motor functions for patients and can benefit the general population by providing a direct link of the brain with robotics or other devices. In this work, we developed a paradigm using deep learning (DL)-based decoders for continuous control of a BCI system and demonstrated its capabilities through extensive online experiments. We also investigate how DL performance is affected by varying amounts of training data and collected more than 150 hours of BCI data that can be used to train new models. The results of this study provide valuable information for developing future DL-based BCI decoders which can improve performance and help bring BCIs closer to practical applications and wide-spread use.

## Introduction

Brain-computer interfaces (BCIs) are devices that allow users to interact with computers or other devices by determining their intentions directly from their recorded brain signals. These fast-growing devices have the potential to help patients suffering from motor or speech disorders by allowing them to control devices such as prosthetic limbs or perform everyday tasks like sending emails or turning on lights. Successful BCIs also have the potential to improve the quality of lives of healthy individuals by allowing them to interact directly with electronics such as cell phones or computers without the need for muscle activation or speech.

The brain recording methods used for BCIs can be generally split into two main categories based on whether they require an invasive procedure or not. Invasive BCIs can often get closer to the source of the brain activity and generally offer good quality signals. Non-invasive brain recording methods such as electroencephalography (EEG) record neural signals from outside the brain and do not require any invasive procedures. Although they carry less risk, the signals from these methods can be limited in signal-to-noise ratio and spatial resolution compared to invasive recording techniques.

BCI applications that require the best available signal quality may be able to justify the medical procedures required for invasive BCIs, but so far invasive BCIs have only been implanted in several dozens of patients. For other applications, non-invasive BCIs provide a safer and more widely available alternative. Among non-invasive methods, EEG is particularly well suited for BCI systems due to its high temporal resolution, portability, and relatively low cost (1). An example of an EEG-BCI setup is shown in Figure 1A. These systems have been used to control devices such as robotic arms (2–4), wheelchairs (5–7), and a virtual (8, 9) or even physical quadcopter drone (10) and could one day be used to control advanced devices like lower limb prostheses (11). Although these EEG-based BCIs have been studied for several decades (12), recent advances in hardware, software, and available computing power have opened up new possibilities for this technology.

**Figure 1.**
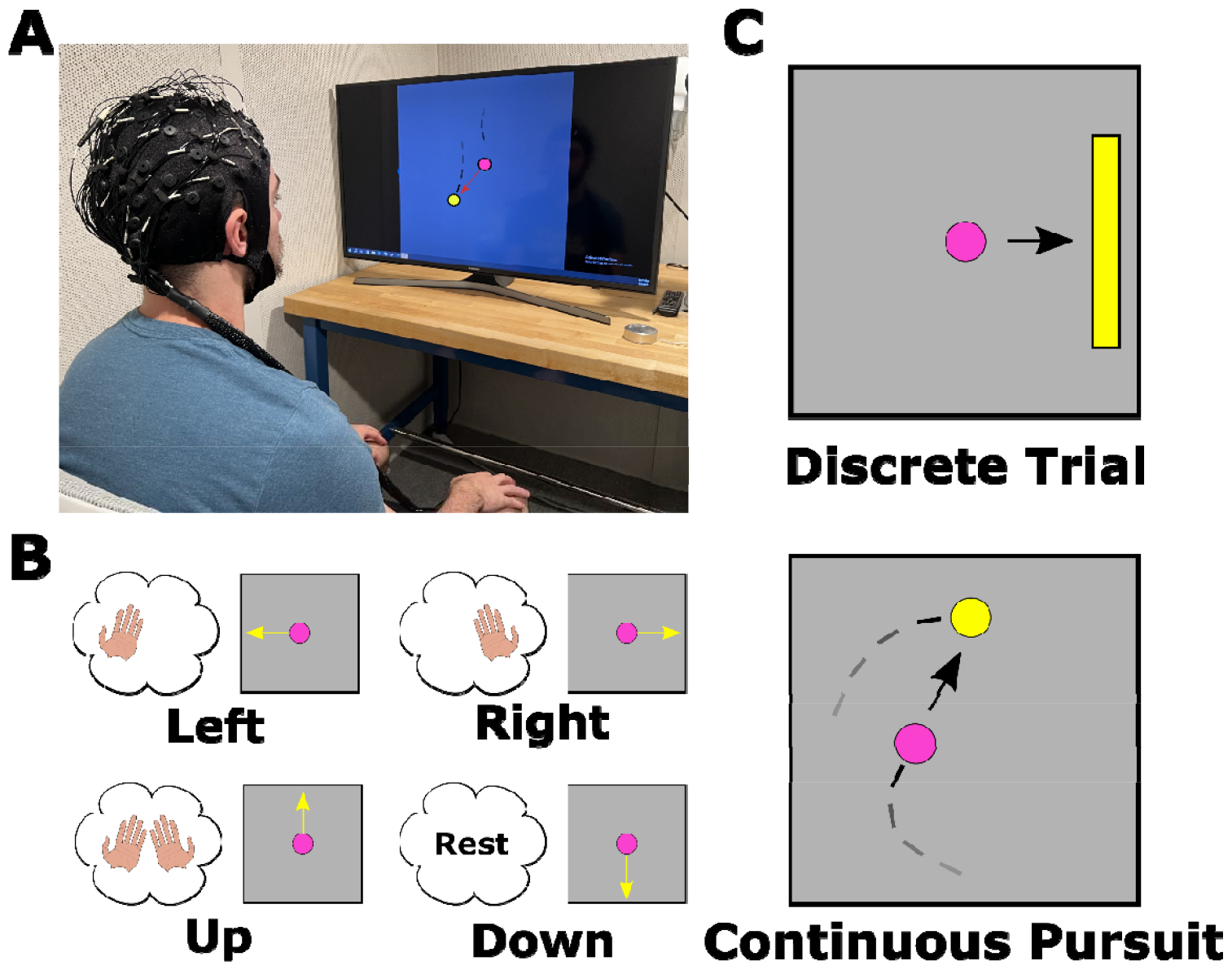
Schematic illustration of EEG BCI. A) A BCI setup, where the user is interacting with a computer with their intention recorded and decoded from EEG. B) An example MI control paradigm for BCI. C) Illustrations of the DT and CP BCI tasks.

One of the common ways to control an EEG-based BCI is the Motor Imagery (MI) paradigm. To perform MI, the user imagines the feeling or sensation of moving a body part, without executing the movement (13). Since this method does not require muscle activation, it can be used by both healthy individuals and patients with some motor impairments (14). In addition, previous studies have shown that MI-based BCIs may be able to help the rehabilitation process for stroke-affected patients (15–17).

At rest, healthy individuals have neural activity in the sensorimotor area of their cortex that oscillates at around 8-13 Hz. These oscillations are typically referred to as the mu or alpha rhythm, and are likely caused by synchronized activity of neurons in the sensorimotor area. When users perform or imagine movements, they reduce this synchronized activity and the amplitude of these oscillations decreases in what is known as Event-Related Desynchronization (ERD) (13, 18). Changes in this frequency band can be detected in the EEG signal, which have also been correlated with BOLD fMRI signals (19), and used to infer when users are imagining the movement of their body parts, allowing them to control a BCI device.

Traditionally, BCI experiments have involved subjects performing MI, or another control paradigm, to produce a specified task or output. In one common setup, subjects are shown a target on one side of a screen and are asked to perform MI to move a virtual cursor to hit the target using predetermined controls (Figure 1B and 1C) (20–23). These experiments, which we will refer to as Discrete Trial or DT tasks, have allowed subjects to achieve effective control over BCI systems as they are intuitive, and users only have to focus on one target outcome in each trial. However, while subjects can obtain good performance in DT tasks, these systems can often take several seconds to produce a single output and may not be suitable for real-world applications that require dynamic control.

A new type of BCI task, Continuous Pursuit (CP), has been recently explored for EEG-based BCIs using MI (2) or actual movement of a limb (24). In this task, subjects are tasked with continuously tracking a target that randomly moves throughout a trial, instead of having only a single target movement (Figure 1C). This task is more complex than similar DT tasks, since subjects need to move their cursor in many different directions throughout a single trial to get closer to the randomly moving target, instead of only having to move in a single direction each trial. Although it is more difficult, this task requires dynamic and continuous control, bringing it closer to the requirements of real-world applications like controlling robotic devices.

There are several potential ways for the subject to control a CP BCI system. In this study, we focus on one paradigm, where subjects interact with the BCI by imagining moving their left hand to move a cursor leftwards, their right hand to move right, both hands simultaneously to move upwards, and neither hand to go down. Collectively, subjects control the movement of the cursor to follow the movement of a target.

One of the other major advances in BCI technology in recent years is the development of more complex decoders. In particular, deep learning (DL) models have been applied to BCI decoding with some success and have been shown to improve performance beyond the abilities of conventional decoders (25–31). This technology has seen rapid growth due to successful applications in many other fields that require complex modelling, such as computer vision and machine translation (32). These models can extract features and model relationships from large and complex datasets, making them an excellent candidate for decoding EEG signals and particularly well suited for taking advantage of subject-specific features. The success of these decoders has been generating increasing interest from the field, and many different DL architectures and paradigms have been recently proposed for BCI decoding (25).

While DL models have the potential to improve BCI performance, there are also several challenges that need to be addressed compared to conventional decoders. Training DL models can require large amounts of training data to achieve better results (26). In other fields that DL has been used successfully, such as image processing, training data is relatively easy to acquire or is already available in large amounts (33). However, BCI data is costly and time consuming to collect and data can be collected with different parameters that may not be compatible such as different sampling frequencies or number of EEG channels (34, 35). In addition, large variances in BCI features between subjects or even different sessions from the same subject can lead to overfitting problems when models are trained extensively on limited data.

Even with these challenges, the potential performance improvements offered by DL-based decoders make them a promising signal processing method for future BCI systems. In addition, methods are being developed to overcome these challenges and take advantage of the strengths of DL. Subject-specific models are commonly trained to reduce the effects of inter-subject variability. Transfer learning is a method used in other fields where DL has been applied to reduce the amount of training data needed for effective models (36). This method, which can involve training at least part of a model on data from a different but related task, has seen increasing interest from the BCI field as DL becomes more popular (37–40). With these methods, and the ability to extract features from complex data like EEG signals, DL-based decoding has the potential to improve BCI systems beyond conventional methods, especially in complex tasks like CP.

In this work, we aimed to improve BCI performance in the CP task by implementing DL-based BCI decoders for online experiments. To accomplish this, we developed a system to label CP data for supervised learning and used it to train several DL models for BCI decoding. We then evaluated decoders based on two different DL architectures in online BCI experiments over several sessions and studied the effects of increasing amounts of training data. We also evaluated additional decoders that used methods that could potentially improve performance even further such as transfer learning, to potentially reduce the amount of training data needed, and recalibration, to reduce the effects of inter-session variability.

Overall, the CP task is challenging and requires both continuous and dynamic control of the

BCI system. However, improving performance in this task through advanced decoding methods like DL can help bring non-invasive BCI devices closer to real-world applications like continuous control of robotic devices. In turn, these advances have the potential to improve the quality of lives of both healthy individuals and motor-impaired patients by offering a direct line of communication with their electronic devices.

## Results

There are several different methods that can be used to quantify BCI performance in the CP task. The primary metric for online experiments is the Mean Squared Error (MSE) between the cursor and the target. To make this number more intuitive, we use the normalized MSE (NMSE) throughout this manuscript. The normalized MSE is the squared distance between the cursor and target, divided by the diagonal length of the square task area. Using this metric, the best performance is 0, where the cursor and target are in the same spot, and the worst possible performance is 1, where they are in opposite corners.

Boxplots are constructed as follows: the center line is the 50th percentile of the data (median), while the bottom and top edges are the 25th and 75th percentiles, respectively. Whiskers are drawn to the furthest data point from each edge that is less than 1.5 times the interquartile range (75th percentile – 25th percentile, or the length of the box). All other points are shown individually as outliers. In each figure, statistical significance is shown by *: p<0.05, **: p < 0.01, and ***: p < Numerical p-values for each test can be found in the Supplementary Materials.

### A. Group-level Results

Figure 2 shows the group level results in terms of cursor-target position NMSE. Figure 2A shows the overall results throughout each session of the experiment. The Traditional (non-DL) decoder performs better than both DL models in the first session, but each of the DL models improves throughout the experiment. Both the Traditional decoder and the Chance runs do not have a clear trend over sessions.

**Figure 2.**
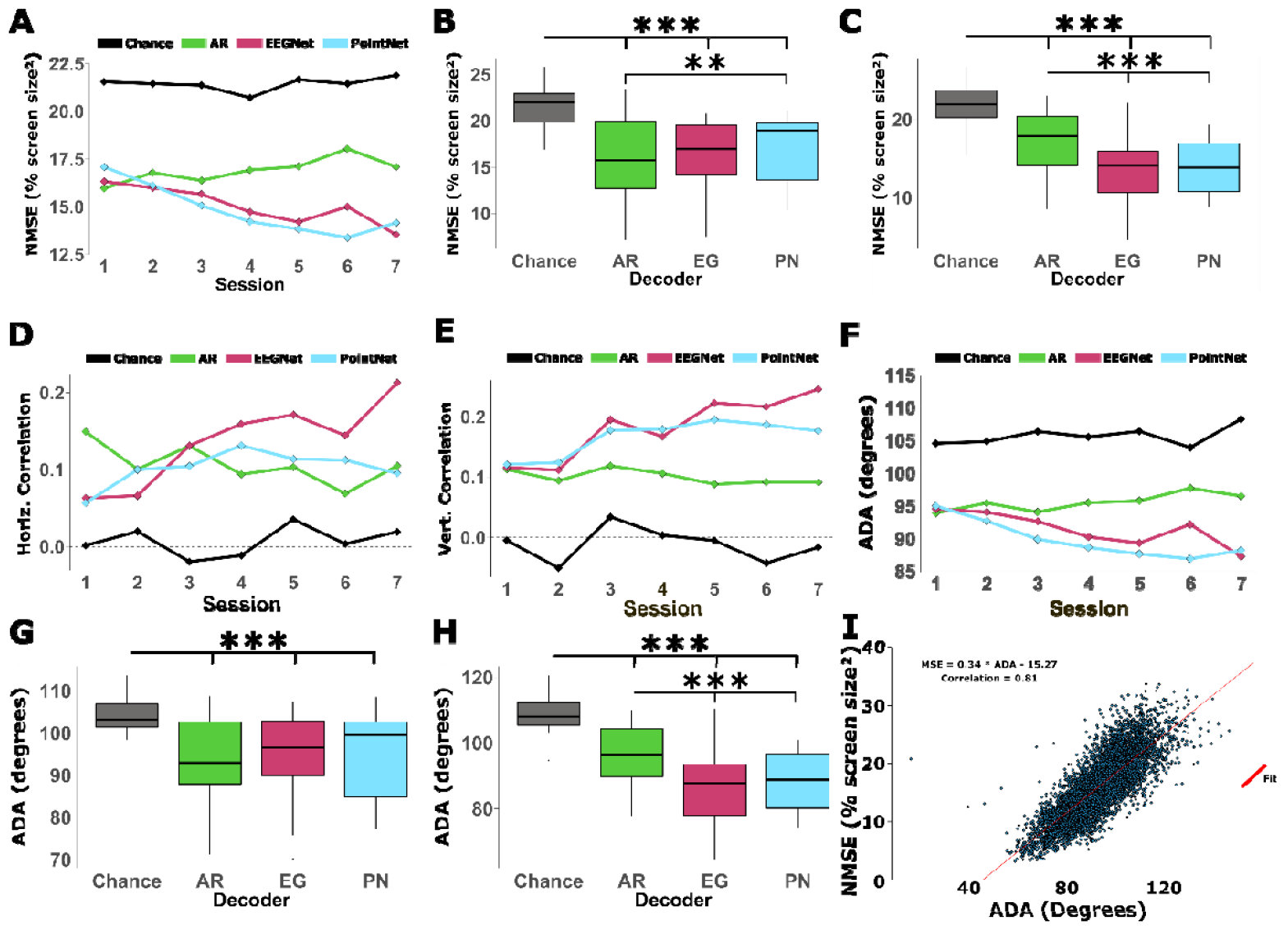
Results of DL decoding in online CP experiments. A) Decoder performance throughout sessions of the study in terms of Normalized Mean Square Error (NMSE). A lower NMSE indicates better performance. B) NMSE performance of decoders in the first session. P-values: ANOVA <0.0001, Chance/AR <0.0001, Chance/EG <0.0001, Chance/PN <0.0001, AR/EG 0.7019, AR/PN 0.0052, EG/PN 0.1134. C) NMSE performance of decoders in the last session (Session 7). P-values: ANOVA <0.0001, Chance/AR <0.0001, Chance/EG <0.0001, Chance/PN <0.0001, AR/EG <0.0001, AR/PN <0.0001, EG/PN 0.2768. D) Horizontal correlation between cursor and target positions (Pearson Correlation Coefficients) over sessions throughout the study. E) Vertical correlation throughout the study. F) Performance of each decoder over sessions in terms of ADA. G) ADA performance in the first session. P-values: ANOVA <0.0001, Chance/AR <0.0001, Chance/EG <0.0001, Chance/PN <0.0001, AR/EG 0.8328, AR/PN 0.4515, EG/PN 0.9216. H) ADA performance in the last session (Session 7). P-values: ANOVA <0.0001, Chance/AR <0.0001, Chance/EG <0.0001, Chance/PN <0.0001, AR/EG <0.0001, AR/PN <0.0001, EG/PN 0.6182. I) Scatter plot of each trial in terms of ADA vs. NMSE. A strong correlation was found between the two and a linear fit is shown in red. Boxplots are constructed in the style of Tukey, showing the median, first and third quartiles, and whiskers up to 1.5 times the interquartile range.

Panels B and C of Figure 2 show the results from the first and last sessions individually. In Session 1, all three decoders perform better than chance level, and the Traditional decoder outperforms PointNet but not EEGNet. In the final session, Session 7, both DL models outperformed the Traditional decoder and there was no significant difference between EEGNet and PointNet. EEGNet improved by 0.43 per session (%screen size^2^) while PointNet improved by 0.56. Supplementary Videos (SV) 1, 3, and 5 show reconstructed trials from this sub-study of two different subjects using EEGNet, and one of the subjects using the PointNet architecture.

Another way to evaluate the performance in the CP task is through the correlation between the cursor and target position. This metric measures the relationship between the cursor and target positions and will be higher when they tend to be in similar locations of the task area throughout the trial. A correlation close to zero implies that there is no substantial relationship between the cursor and target movements, making this an intuitive benchmark for a random baseline performance. As expected, the experimental chance level results fluctuate around zero correlation in both the horizontal and vertical directions.

The correlation results were split into each axis, with the correlation between horizontal cursor and target positions shown in Figure 2D, and the vertical results in Figure 2E. In both plots, the traditional decoder remains relatively level across the sessions, while both DL decoders improve over time. In both cases, the EEGNet performance reaches a higher performance than PointNet by the last session, and seems to be continuing on an upward trend while PointNet starts to plateau over the final few sessions.

Figure 2F illustrates the group-level CP results in terms of the Average Difference Angle (ADA) metric over each of the sessions (see Materials and Methods for details on this metric). Here, all three of the decoders tested in this study perform similarly in Session 1, but both DL models improve over time and outperform the traditional decoder by the last session. The results specific to Session 1 and Session 7 are shown in Figures 2G and 2H respectively.

We chose to use ADA for training and evaluating the DL models offline since this metric can be obtained without simulating a cursor path in an offline CP trial. However, the cursor-target NMSE remains the primary metric for quantifying online performance and may be more intuitive than ADA. To study how well ADA may substitute for NMSE in offline analysis, we looked at the correlation between the two metrics. Figure 2I shows a scatter plot of each online CP trial comparing the distance and ADA results, along with a linear fit. The result over all online trials was a correlation of 0.81 between distance and ADA, showing a strong relationship between the two metrics.

### B. Cursor-Target Lag

During the CP task, BCI decoders can suffer from the delay period between when the cursor-target direction changes and when the decoder can update accordingly. This delay period can have several contributing factors including: the subject’s reaction time and ability to change their motor imagery quickly, as well as the ability of the decoder to detect changes in motor imagery features and modify its output in a timely manner. While this lag period can be a challenge for the CP task, it may not have a substantial effect on traditional BCI tasks that have a single target in each trial, like the DT paradigm.

To investigate this delay, or lag, between changing target directions and decoder output, we shifted the cursor positions over time and calculated the cursor-target distance at each lag. The lag that produced the best performance (lowest cursor-target NMSE) was recorded for each trial. Figure 3A shows a histogram of the results over all the decoders. The large spike after a few seconds shows that the performance in most trials is better using the target position from a few seconds in the past, rather than the current target position. Figures 3B, 3C, 3D, and 3E show the same histogram for the experimental ‘chance’ runs, the traditional model, EEGNet, and PointNet, respectively. Similar spikes around 1.5 seconds are present in each of the decoders, while the chance level runs do not have any spike, showing that the delay is not present in runs without any BCI control.

**Figure 3.**
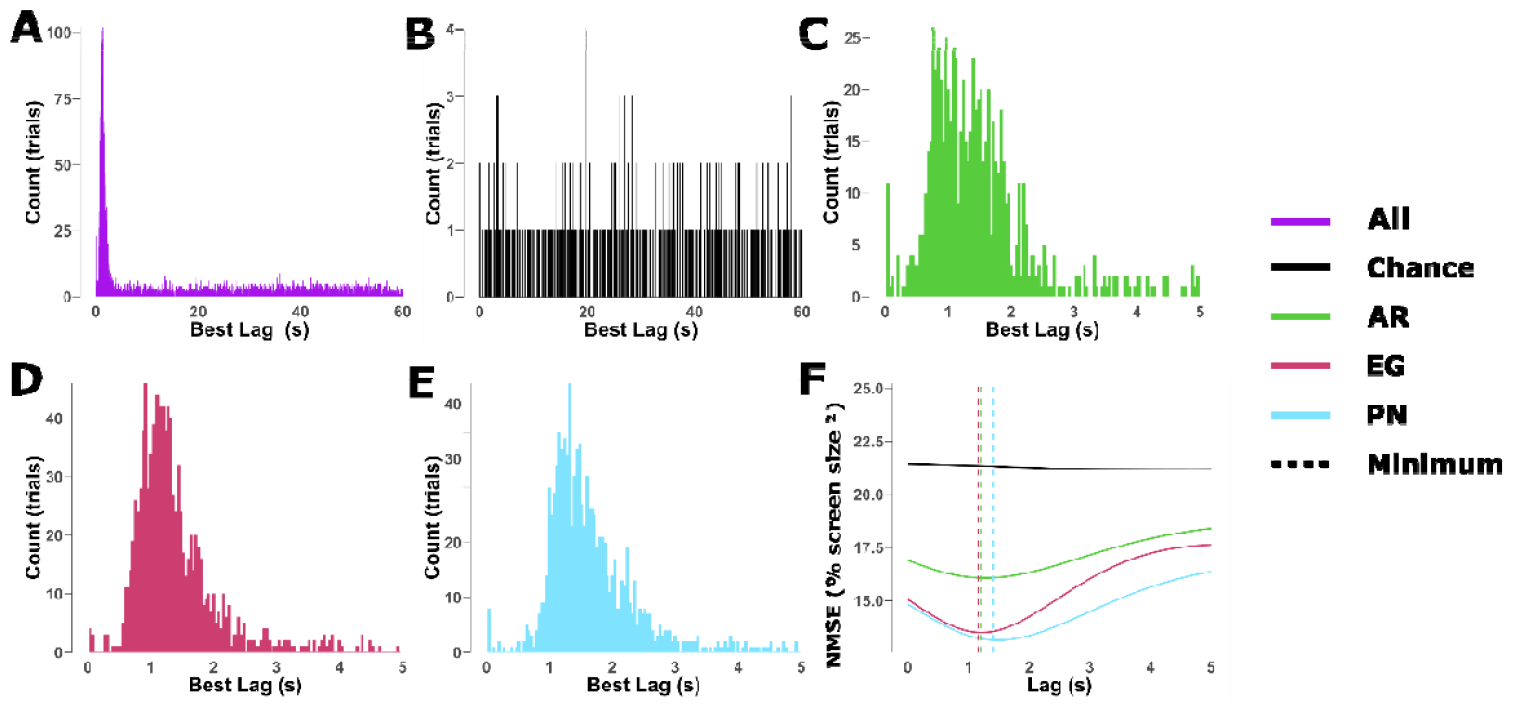
Cursor position lag. A-E) Histograms of the cursor position lag that results in the best performance for each trial. The large spikes around 1-2 seconds suggest that there is a short delay between a change in target direction and the decoder output. F) CP performance in cursortarget NMSE for each lag duration. The low points in each curve show the estimated BCI performance from removing the delay.

Figure 3F shows the grand-average NMSE over all sessions for each decoder as a function of lag. The performance of each BCI decoder improves with the delay, up to a peak between 1-2 seconds, while the ‘chance’ level performance remains steady throughout any lag. The traditional model performed best after a lag of 1.28 seconds and the NMSE was improved by 0.8 (%screen size^2^). EEGNet’s best lag was 1.2 seconds and improved by 1.6, while PointNet was later at 1.48 seconds and improved by 1.7. Supplementary Videos 2, 4, and 6 show the same trials from SV1, SV3, and SV5, but with the lag adjusted for by shifting the cursor position forward in time.

### C. Transfer Learning and Recalibration

One of the main disadvantages of DL-based decoders is that they can require a larger amount of training data to achieve optimal performance compared to other methods. Transfer learning (TL) is a method used in some DL applications that could potentially reduce the amount of training data needed for these models by pre-training them on data from previously studied subjects. We hypothesized that pre-training DL-based decoders on the data from the first part of this study could help the resulting TL-based models perform well in early sessions for new subjects, even with the limited subject-specific training data available in the first few sessions.

In addition to needing enough training data, DL-based decoders are also prone to overfitting on the training dataset. The large variances in BCI data between subjects, and even between sessions for the same subject, may cause reduced performance when models are fit too heavily on training data that is from a different subject or day than the online testing session. To help models adapt to the current task, we hypothesized that recalibrating the models using data from the current testing session could help improve performance.

We compared both TL and recalibration (CL) to the same EEGNet paradigm we used in the first sub-study (DL). Like the first part of the study, the DL model requires subject-specific training data and could not be used until Session 2. We chose to include 4 runs from the traditional decoder in Session 1 to compare with the TL decoder in the first session, where the TL model has not been trained on any subject-specific data.

The results over the first 4 sessions are shown in Fig. 4A. All three models performed similarly at the group level, and both TL and DL saw improvements each session from Session 2-4. Interestingly, the TL models did not improve from Session 1 to Session 2 at the group level (average across all subjects). Both the TL and DL training methods achieved similar performances throughout Session 2-4.

**Figure 4.**
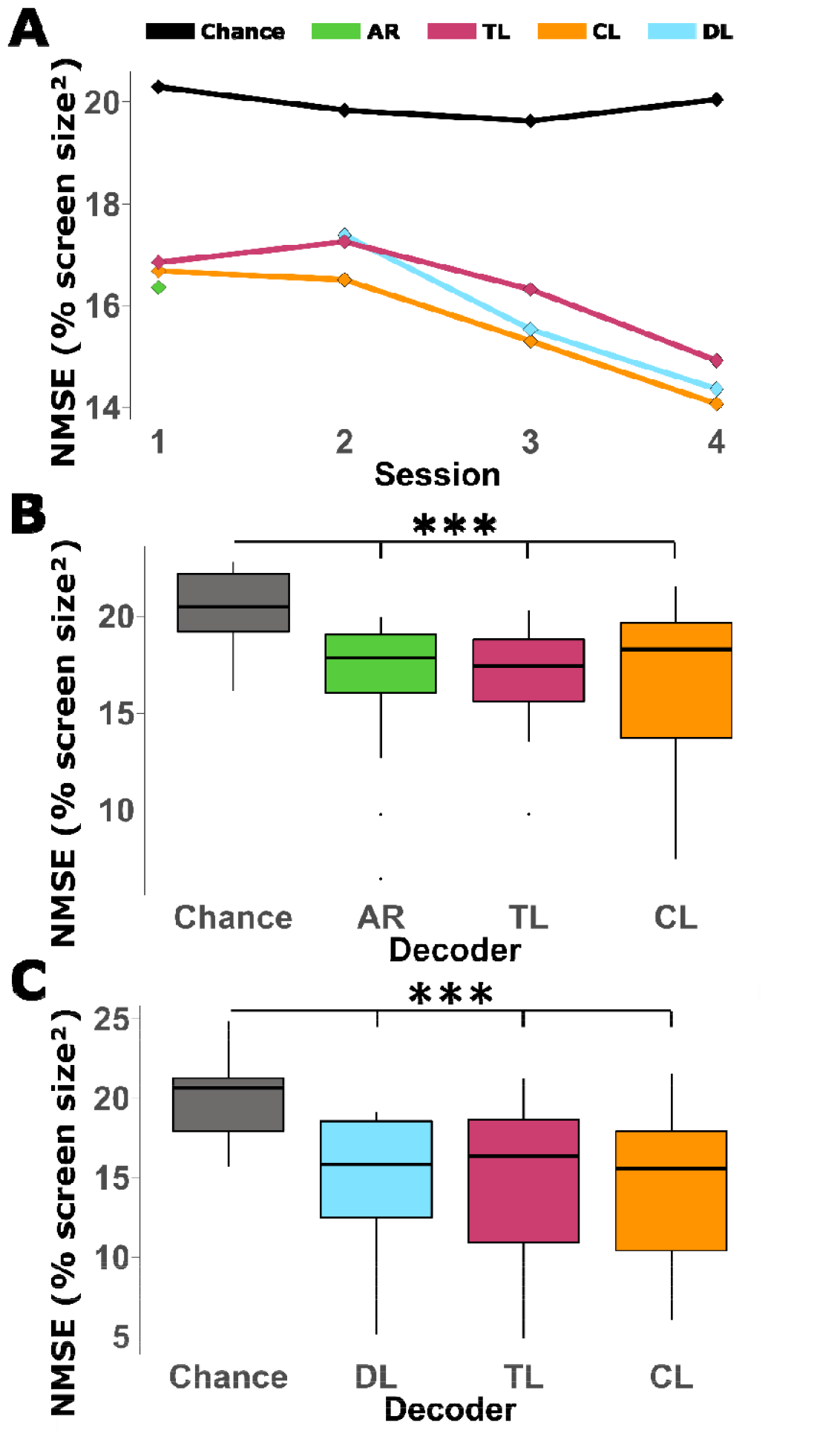
Transfer learning results. A) Overall results from sessions 1-4 in terms of NMSE B) Results from the first CP session, Session 1. P-values: ANOVA <0.0001, Chance/AR <0.0001, Chance/CL <0.0001, Chance/TL <0.0001, AR/TL 0.6224, AR/CL 0.8597, TL/CL 0.9753. C) Results from Session 4 of the CP experiments. P-values: ANOVA <0.0001, Chance/DL <0.0001, Chance/CL <0.0001, Chance/TL <0.0001, DL/TL 0.3723, DL/CL 0.8331, TL/CL 0.0676. Boxplots are constructed in the style of Tukey, showing the median, first and third quartiles, and whiskers up to 1.5 times the interquartile range.

Looking at the results from Session 1 (Fig. 4B), the traditional decoder and TL models reach similar performance, and the CL models had a similar median but larger variance. In the final session (Fig. 4C) had similar results where each of the three training methods achieved similar performances, but CL had a large variance with some subjects benefiting substantially.

Lastly, we compared the performance of the first sub-study with the faster speed to the second sub-study using the slower speeds (Supplementary Fig. S1). For this analysis, we compared only the runs using EEGNet without TL since this was the only condition present in both sub-studies. Although we lowered the cursor and target speed to make the task easier for subjects, the results show that the performance using both speeds was not substantially different.

Furthermore, we looked at the effects of lag on both speeds (Supplementary Fig. S2) and found that the lag affected performance more using the faster speed. Although the slower speed performance was less affected, the optimal lag was longer than the faster settings (2.04s vs. 1.34s). This may be because the slower cursor speed required more time for subjects to correct their course.

## Discussion

In this study, we designed a new framework to train deep learning-based EEG decoders for the continuous pursuit (CP) task and evaluated the performance of two different DL architectures and a traditional decoder over seven BCI sessions. We found that both DL-based decoders improved throughout the study and outperformed the traditional decoder by the final session. Both the EEGNet- and PointNet-based decoders performed similarly across several metrics including cursor-target distance, correlation, and ADA. When we implemented a transfer learning technique by pre-training DL-based decoders on data from previous subjects, we found that the pre-training did not significantly improve BCI performance in early sessions compared to training from scratch. However, recalibrating models by training on small data samples from the current online session helped to improve outcomes in some situations. Overall, using DL-based decoders improved their performance throughout the course of the study and outperformed the traditional decoder once enough training data became available.

Motor imagery (MI) BCIs aim to help individuals to control an external device without the use of conventional muscular activity. In the current study, the MI of left hand, right hand, both hands or none was used to accomplish online continuous control of a virtual cursor in a 2D space. Despite the relatively simple MI paradigm, this reflects the state of the art of non-invasive BCI, constructing a mapping relationship between BCI controlled tasks and the intended outcome. Using DL-based decoding in online experiments, we demonstrated in this work that the proposed BCI system is able to control a virtual object to move continuously with changing direction and may be used as a neuroassistive device. Further developments may translate such a CP paradigm for more advanced applications, such as continuous control of a robotic device or the use of a more complex MI paradigm. In addition, prior study suggested the merits of MI BCI to help motor recovery in stroke patients using a simple discrete MI task (41), it would be interesting to test if CP MI tasks would further help stroke rehabilitation given the complexity of CP through motor imagery.

The CP task is an extension of more traditional discrete trial tasks such as in: (21, 23). While these tasks require the user to continuously control a cursor to hit a target, the target is only in a single location in each trial and therefore only requires moving in a single direction. Continuous tracking paradigms such as CP have the target randomly move around the workspace, requiring subjects to change the cursor’s direction to follow the target throughout the trial. Edelman et al. (2) developed the CP paradigm used in this study by having subjects use hand MI to move a virtual cursor and/or robotic arm to continuously pursue a randomly moving target, and showed that this could be achieved using MI with good performance with the aid of EEG source imaging. In this work, we demonstrate that using EEG sensor space data with the aid of DL-based decoders, humans are able to track randomly moving targets with good performance, by means of the MI paradigm.

More recently, Mondini et al. (24) used a trajectory reconstruction paradigm to decode subjects’ executed hand movements from EEG, and were able to move a robotic arm to continuously track a randomly moving virtual snake. In this work, we demonstrate that without explicit hand/arm movement, users can perform CP tasks using MI EEG signals. Applying DL-based decoding to the online CP task can help improve performance in this difficult task and similar models could potentially be trained for other paradigms in the future, such as executed or imagined hand kinematics.

While DL-based decoders have been implemented for other BCI tasks, the CP paradigm lacks a clear label for the data to use for supervised learning purposes. This makes training DL models using the conventional methods more difficult compared to discrete trial BCI tasks, where there is a single target for each trial and therefore a clear label. To solve this problem, we developed a CP task labelling method, described in the Materials and Methods section, that uses the vector between the cursor and target positions, as a label. Similar techniques have been used in the invasive BCI field to allow decoders to adapt to users’ intentions throughout trials with the set target in each trial in DT tasks like center-out and center-out-and-back (42, 43). The labelling system used here expands to the CP paradigm, where both the cursor and target move throughout the trial and are continuously changing. By using the cursor-target vector as a label for data in the CP paradigm, we are able to use methods that rely on supervised learning, such as DL, for the CP task which were previously unavailable.

Although this paradigm allows us to apply a label to the CP data for supervised learning, it also creates two potential problems for the decoders. First, the labelling system assumes that the subject is always trying to move in the optimal direction towards the target. Although this is the best direction for the cursor to get closer to the target, subjects may not always be trying to move in this direction during the actual experiment, which could generate inaccurate labels. Second, the MI paradigm that the subjects were instructed to move only consisted of four cardinal directions (left, right, up, and down). To move in other directions, the subjects had to perform a combination of these MI tasks, and it was up to each subject to determine their best strategy for combining the movements. How subjects chose to integrate these movements could make the difference between producing decodable features for multiple directions, or just the four cardinal directions.

Even with these challenges, this work shows that the models were able to capture some relationship between the label and subjects’ EEG recordings and provides evidence that the labelling system is suitable for training supervised learning models for the CP task.

The overall group results in Figure 2 show the development of each decoder throughout the study sessions. In the first session, all three decoders perform better than chance level, showing that the DL-based decoders can perform even when trained using only a single session of subject-specific data. However, the traditional AR-based decoder had the best average performance and statistically outperformed the PointNet model. Throughout the study, both DL-based decoders improved when given increasing amounts of training data and outperform the traditional decoder by the final session (Fig. 2C). Overall, these results show that DL-based decoders improve when given increasing amounts of training data and may even keep improving past the 7 sessions studied here.

We originally chose to implement two DL architectures to show that this paradigm could accommodate different models and to examine any performance differences between architectures. EEGNet was chosen since it is an established decoder for EEG and BCI applications while we chose to implement the PointNet model to test a new architecture for BCI that has seen success in other fields. The results show that both models performed similarly, so the additional complexity of PointNet may not be justified in this case. Still, the architecture may be useful for other BCI applications, especially where the input can be represented as a group of points in a multi-dimensional space or point cloud. For example, EEG source imaging based BCI (2, 44, 45) can estimate brain activity from inputs of scalp recordings by electrodes distributed in three-dimensional space which could be represented as a point cloud and may be a potential application for the PointNet architecture.

Interestingly, the performance of the traditional decoder seemed to stay consistent throughout the study instead of improving as subjects got more experienced. One reason that we did not see an improvement in performance due to learning may be that all the subjects recruited for this study were experienced and had already participated in at least 2 BCI sessions prior to this experiment. This means that any performance improvements from learning in sessions that subjects completed before this study are not shown in the results here.

Another potential factor could be that subjects may be able to tell the difference between the three decoders during the experiment based on the cursor movement and their observed performance. Subjects were made aware that different decoders may be used throughout the experiment but were not told which decoder would be used for each trial. Differences in decoder performance may have caused some confusion or conflict for subjects when the BCI system started reacting differently to their intentions. Although we believe that testing multiple decoders in each session was essential to this paradigm, this could have affected the results in some cases. For example, this may be one explanation for the slight decrease in performance for the traditional decoder towards the final sessions. As the DL decoder performance improved with more training data, subjects may have noticed the large performance difference during the AR trials and may have been discouraged during these trials where it was more difficult to control the cursor. Although subjects were instructed to try their best during every trial, lower motivation during the more difficult AR trials could lead to lower performance.

Since the cursor-target NMSE is the primary metric for the CP task but cannot be readily used offline, it is important to determine if ADA can be a used as an offline approximation of NMSE. To study this, we looked at the correlation between cursor-target NMSE and ADA shown in Figure 2I. The high correlation (0.81) shows that both ADA and distance follow a similar trend and that a low ADA offline suggests that a similar online setup would achieve a good NMSE. However, there is still a large variance off the linear trend, meaning that ADA is not a perfect predictor for online performance.

During the experiments with the faster speed, we noticed a substantial lag between the target position and cursor movement. We hypothesized that this lag could be caused by both subject reaction times and the decoder’s ability to change its output. The results in Figure 3 show an analysis of shifting the cursor position vector in time. The histograms in panels A-E show how frequently each time-shift resulted in the best cursor-target distance for each trial. The large peaks around 1-2 seconds for each BCI decoder show that the NMSE improves by shifting the cursor position forward 1-2 seconds in time suggesting that the cursor position lags the target by about 1-2 seconds. Figure 3F shows the average cursor-target NMSE across trials for each decoder as a function of lag. Here, all three decoders see peak performance after a short delay. The fact that all three decoders have similar lags suggests that this delay may be primarily due to the CP task or MI paradigm, instead of the individual decoder designs. The time it takes subjects to react to changes in the cursor and target positions and adjust their MI intentions accordingly will add a delay between the cursor and target movement, regardless of the BCI decoders used. This lag between target movement and decoder output presents a unique challenge for the CP paradigm. While real-time and dynamic control is desired for continuous tasks, the subjects’ reaction time and the internal buffers of decoders could limit the ability of BCI systems to react quickly. This is even more pronounced in higher speed CP tasks, where the subject is likely to have to change their intention more often and decoders will have less time to modify their output.

One of the primary drawbacks of DL-based decoders is that they can require large amounts of training data before they can perform well. This issue is especially relevant for BCI tasks, where collecting data can be costly in both time and money. In this study, we implemented a TL paradigm to reduce the amount of training data needed by using data from the 14 subjects studied in the first part of this work to pre-train DL models for new subjects. In addition, we attempted to further improve performance for these new subjects by fine-tuning models mid-way through each session using data collected from the current session. Since we hypothesized that the TL models would outperform the standard DL training paradigm, we performed this recalibration step on the TL models.

The overall results from this second set of experiments (Figure 4A) follow a similar trend to the original experiments. Each of the DL-based decoders improved in successive sessions as more subject-specific training data became available. Overall, the TL and DL models reached similar performance in sessions 2-4, showing that our specific implementation of transfer learning did not benefit performance in early sessions. Interestingly, the TL decoders reached a similar performance to the traditional decoder in the first session (Figure 4B), when no subject-specific data were available. Since the TL models had no subject-specific training data at this point, it makes sense that they would not be able to take advantage of any subject-specific features, which could be one explanation for the similar performance. Both these results suggest that the main benefits of DL-based decoding lie in the models’ ability to extract subject-specific features.

For this study, we implemented a basic transfer learning method involving pre-training a model on existing subject data, then fine-tuning the model on new subjects. While this method allowed us to initially explore the effects of TL and demonstrate the implementation in an online CP paradigm, there are a variety of more sophisticated TL algorithms developed for BCI applications that may be more effective (38). We believe this work demonstrates the potential for TL to improve complex online BCI applications like CP and that the data collected from this study may be helpful for testing more advanced TL methods that can reduce the amount of training data needed for new subjects or even improve performance overall.

The results of same-session recalibration are interesting, but not conclusive. The recalibrated models reached a better average performance than the TL models that they are based upon in each session after the first, but do not reach statistical significance. Our current implementation of recalibration uses only data from four runs to update the models mid-session, and we do not have any way to choose the best model weights among the five training epochs. Future iterations of this method could include a validation set or use a different way to update the model weights, such as a different learning rate or only updating specific layers. The recalibration results shown here suggest that some method of updating the models mid-session may help to reduce intersession variance and warrant further investigation.

This study demonstrates the potential of using DL-based decoders for online BCI decoding in challenging tasks and shows that subjects can achieve strong performance with these models. However, additional work could be done to further improve performance and help optimize the training process. In this study, we compared the results from three different decoders: EEGNet, a well-established DL architecture for EEG BCIs, PointNet, our exploration into a new DL architecture, and the AR decoder, a traditional decoder used in BCI research. There are many other algorithms that have been developed for BCI decoding, including other DL architectures and non-DL methods that are more advanced than the AR model used here. Future work should be done to implement these models for online CP experiments, and rigorously compare the resulting performances to identify the best available decoders. Although the models performed similarly in this case, other DL architectures could be designed specifically for the CP task and could produce better outcomes. The experiments from this work produced over 150 hours of CP BCI recordings that could be used as a rich training dataset for developing such models. We hope that the work done here will enable and encourage more studies towards developing better decoding algorithms for complex BCI tasks like the CP paradigm.

In both parts of this study, the DL model performance is still improving between the second to last and final session studied, suggesting that the models could still benefit from additional training data. A long-term study with many sessions for each subject could help to further characterize the relationship between DL-based decoder performance and training data and may be able to determine the full potential of these models.

Lastly, transfer learning and recalibration were studied briefly here, but each method could warrant a study of its own. Although our implementation of TL did not seem to benefit the DL-based decoders, transfer learning has a strong foundation and has seen success in other fields and even some BCI applications (37–39). The recalibration method here did improve performance in some cases but could be limited by the amount of recalibration data available and there may be better ways of fine-tuning the models mid-session. Although this work shows a foundation for implementing DL-based decoding into an online CP BCI system, future work could be done to further improve this setup. Both DL and TL are active research areas, both in and outside of the BCI field, and more advanced decoders or TL algorithms could help improve user performance. In addition, the data collected in this study may be useful for constructing DL and TL models specifically for the CP task.

Applying DL methods to online experiments helped to improve subjects’ performance in the challenging CP task. We found that it was difficult for subjects to precisely follow the target for an entire 60 second trial, but good performing subjects were able to get close to the target and could recover from large errors (see Supplementary Videos for some examples). More work is needed to further improve subjects’ BCI control in complex tasks, but the ability of subjects to repeatedly reach a randomly moving target shows promise that non-invasive BCIs could be used for complex applications in the near future. Even without perfect control, these systems can be enhanced with post-processing and shared control to be useful for real-world tasks, similar to how auto-correction improves spelling applications. In addition, the results presented here are from subjects experienced with BCI control. Future work should be done to investigate the use of online DL decoding for naïve subjects as well as in motor-impaired populations.

In summary, we have developed a novel labelling paradigm and offline performance metric that allowed us to train deep learning (DL)-based decoders for the continuous pursuit (CP) BCI tasks, and rigorously evaluated the DL-based CP BCI in a group of 28 human subjects. Two different DL-based decoders were evaluated over seven BCI sessions and both models were found to improve throughout the study when additional sessions of training data became available and outperformed a traditional BCI decoder after just a few sessions. A transfer learning paradigm and mid-session recalibration technique were also evaluated and show the potential to benefit performance, although more work is needed to optimize these methods. This work introduces online, DL-based decoding to the CP task and shows how these decoders have the potential to improve subjects’ control in continuous BCI systems. Although CP is challenging, strong performance in this task can open the door to more complex and intricate BCI systems with real-time dynamic control and can help move non-invasive BCIs closer to real-world and clinical applications.

## Materials and Methods

In this work, we focused on studying EEG-BCI systems based around the MI paradigm. To control the BCI system with this method, subjects were seated in front of a computer monitor and asked to imagine the feeling or sensation of moving their hands (Figure 1A). Performing hand MI causes ERD in the alpha band of the EEG signal (8-13 Hz, this is sometimes referred to as the mu rhythm, but we will use the term alpha rhythm throughout this manuscript) in the contralateral sensorimotor area, usually around either C3 or C4 (46). This modulation of brain activity can be detected and used to infer the user’s intentions using signal processing methods that we refer to as decoders. The output from these decoders can then be used to control applications such as robotic devices or, in this case, a virtual cursor.

In the CP task, subjects are asked to use MI to control a virtual cursor to move around a 2D screen. They are instructed to imagine moving their left hand to move the cursor to the left, their right hand to move it right, both hands simultaneously to go upwards, and to rest without performing any MI to move the cursor down (Figure 1B). In this paradigm, a virtual circle acts as the target, which randomly drifts around the screen area. The subjects’ goal for the CP task is to move their cursor using the MI controls to track the target as close as possible (Figure 1C).

### A. Subject Recruitment

This study consists of two experimental parts. In the first sub-study, we recruited 15 healthy human subjects (average age: 25.2, 14 right-handed, 8 male) for 8 sessions of CP BCI and fourteen subjects completed all 8 sessions. In the second sub-study, we recruited an additional 15 healthy human subjects (average age: 22.13, 13 right-handed, 8 male) for two BCI training sessions and 4 CP sessions. Fourteen of these subjects completed the 6 total sessions. The subjects in each sub-study were unique and no subject participated in both.

Fliers were placed around Carnegie Mellon University campus and the greater Pittsburgh area to recruit subjects. This study was approved by the Institutional Review Board at Carnegie Mellon University and each subject provided written consent to the protocol before participating. For the first sub-study, all participants were required to have completed at least two previous sessions of MI-based BCI cursor control before beginning the experiment. If a subject was recruited with less than two sessions of experience, they performed discrete trial cursor control BCI sessions to meet this requirement before beginning this study. Experienced subjects were recruited for the first sub-study to reduce any effects of subjects learning MI on the BCI results and to focus on the effects of additional training data on DL-based decoders. In addition, we hypothesized that data collected from when subjects are first learning to perform MI may not be good for training the DL decoders and could reduce overall performance, even in later sessions. For the second sub-study, all subjects recruited had no prior BCI experience, but each subject performed two sessions of discrete trial BCI on different days as part of the study design. Even though we were not focused on the effects of additional training data for this part of the study, we still required subjects to undergo the two training sessions to avoid fine-tuning DL and TL models using the data from when subjects are first learning MI.

### B. Signal Acquisition

EEG signals were recorded using the Neuroscan Quik-Cap with 64 channels and amplified with the SynAmps 2/RT system from Compumedics Neuroscan. The electrode caps were configured in a modified version of the international 10/20 system, with the reference electrode in the center between Cz and CPZ electrodes. At the beginning of each session, all electrode impedances were reduced to below 5 kΩ. Data were collected using the Neuroscan Acquire application at a sampling rate of 1kHz, low-pass filtered under 200Hz with a notch filter at 60Hz to remove potential interference from AC electronic devices.

The BCI2000 program (47) was used to run the BCI experiments and cursor control application. Data packets were streamed from the acquisition software and processed at 40ms intervals for signal processing. The amount of data used to make a classification was different for each decoder, but each decoding method produced an output every 40ms to update the cursor velocity.

### C. Continuous Pursuit Task

The continuous pursuit task was run using custom python scripts made for BCPy2000, part of the BCI2000 program. In this paradigm, the subject is tasked with moving their cursor, displayed on the screen as a circle, in two dimensions to track a virtual target as closely as possible. The subjects used the motor imagery paradigm described above to move the cursor, and each decoder provided two outputs: one for the horizontal axis and another for the vertical.

The virtual target in this task was also a circle with the same size as the cursor. The target was made to randomly drift around the screen in each trial by applying accelerations chosen from a normal distribution every 40 milliseconds. Following the paradigm from later experiments in the original CP study (2), we chose to not allow the cursor or target to wrap around the screen to be consistent with controlling a physical object. To deter the target from remaining on the boundary of the screen for long periods, we applied a repulsion acceleration to the target if it drifted towards the edge of the screen.

### D. DL Training Paradigm

Supervised learning is one of the standard paradigms for training DL models. This method requires the training data to be labelled with corresponding ground truth outputs. For BCI tasks, these labels can be things such as the desired direction when moving a cursor or device, or a selected letter in a speller. BCI tasks with only a single target in each trial, such as DT tasks, have a clear label for each trial and are well-suited for supervised learning. In contrast, the CP task does not have one clear target or direction for each trial that can be used as the label so additional steps are required to use this data for supervised learning.

The subjects’ primary goal in the CP task is to move their cursor as close to the target as possible throughout the trial. Therefore, the optimal direction for them to move their cursor to reduce this distance is the vector from the cursor position to the target position (example shown in Figure 5A). A similar procedure has also been used in invasive BCI literature to help improve decoders during some tasks with the set target in each trial (42, 43). Although this vector may not be the exact direction that the subject is intending to move at any point in time, it should be a close estimate if the subject is attempting the task correctly.

**Figure 5.**
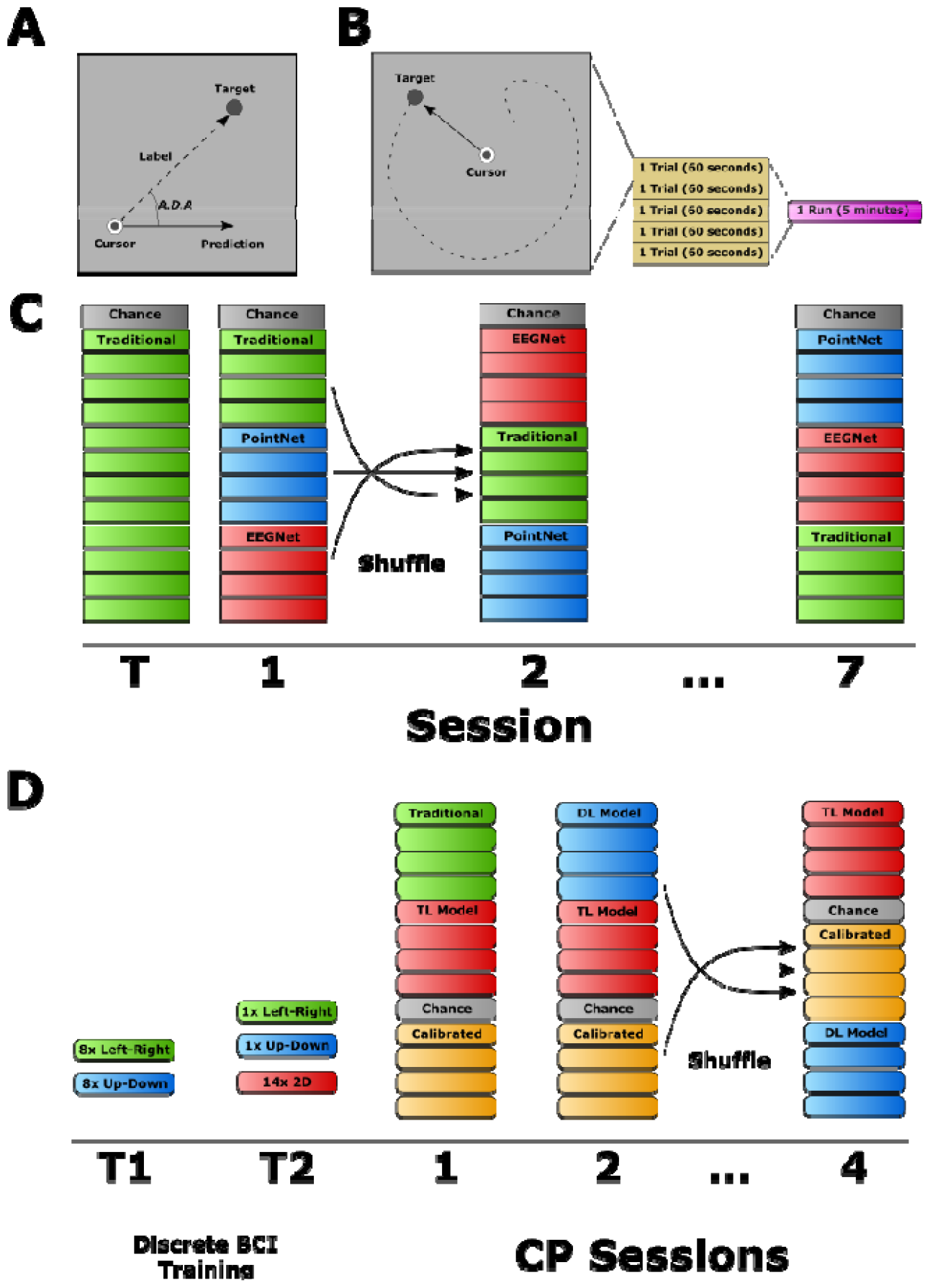
Study Design A) Labeling system for CP data. The vector between the cursor and target positions is used as the label for supervised learning. The angle between a model prediction and true label is used as an offline performance metric: Average Difference Angle (ADA). B) Breakdown of a CP run. Each trial lasts 60 seconds. Each run consists of five trials and lasts about 5 minutes. C) The main study design. Subjects performed eight sessions of BCI experiments, each with 12 BCI runs split between various decoders and 1 additional run to estimate chance level. D) The study design to evaluate transfer learning and recalibration. The setup is like Panel C, but subjects performed two training sessions of DT BCI and four sessions of CP.

The DL models were implemented and trained using custom python scripts and the Pytorch framework, details on model training can be found in the supplemental materials. Preprocessing was different for each DL architecture and more information can be found in the corresponding sections.

In addition to designing a labelling system for the CP task, we also needed to find a way to evaluate CP performance in offline experiments. Cursor-target distance and correlation are informative metrics for evaluating online experiments, where the cursor and target positions are available, but they are more difficult to calculate in offline situations where the positions must be simulated. In an offline task, such as evaluating one set of weights for a DL-based decoder, the outputs from the offline model could be used to simulate a cursor moving, just like in the online task. However, the EEG data are recorded from the online experiment, so when the cursor and target in the offline simulation are in different positions than they were during the online task, the subjects’ intentions during the EEG recordings may not match up with the correct direction for the offline simulation.

To evaluate the DL models’ performance without using the cursor-target positions, we chose to use the angle between the models’ predictions and the labels as a performance metric (Figure 5A). Here, a lower average angle indicates that the prediction is closer to the true label suggesting the model will perform better in an online CP experiment. This method, which we refer to as the Average Difference Angle (ADA), does not rely on cursor or target positions, only the labels which are available from the dataset. This means that the method can be calculated offline without having to simulate a CP trial, and can be used to evaluate DL model performance during training.

### E. Traditional Decoder

To provide a baseline performance for comparing the DL models studied in this work, and to obtain the initial training data for supervised learning, subjects completed several runs of the CP task each session using a traditional online BCI decoder. For this purpose, we chose to use the AR Signal Processing module, part of the standard BCI2000 software. This class of decoder is well established for BCI experiments (21, 48–51) and does not require any training data before use, allowing us to implement it for each subject’s first session. More information about the implementations of the traditional decoder and two DL decoders used here can be found in the Supplementary Materials.

### F. EEGNet

Following the increasing interest in DL-based decoding for BCI, there have been a number of DL architectures developed specifically for EEG BCIs (25, 29). Our previous work compared several of these models on two offline datasets and found that they performed similarly overall, but one model in particular, EEGNet (52), outperformed the others on one of the datasets. Due to EEGNet’s performance in this offline study and well-established use, we chose to implement this model for the online CP experiments.

### G. PointNet

In addition to implementing EEGNet for the CP task, we also chose to test another architecture in online experiments as well. For this purpose, we adapted the PointNet architecture for BCI decoding (53, 54). The PointNet architecture is designed around point cloud representations of the task as inputs to extract spatial patterns or features. In other fields, this point cloud can be a collection of points that make up a 3D scene, but here we used the collection of electrodes and their 3D locations in space (relative to each other) as the point cloud.

Our implementation of the PointNet architecture is shown in Figure 6. The model begins with the frequency spectrum input, which is passed through three successive set abstraction layers, a fully connected layer, batch normalization, a RELU activation function, dropout (40%), and a final fully connected layer.

**Figure 6.**
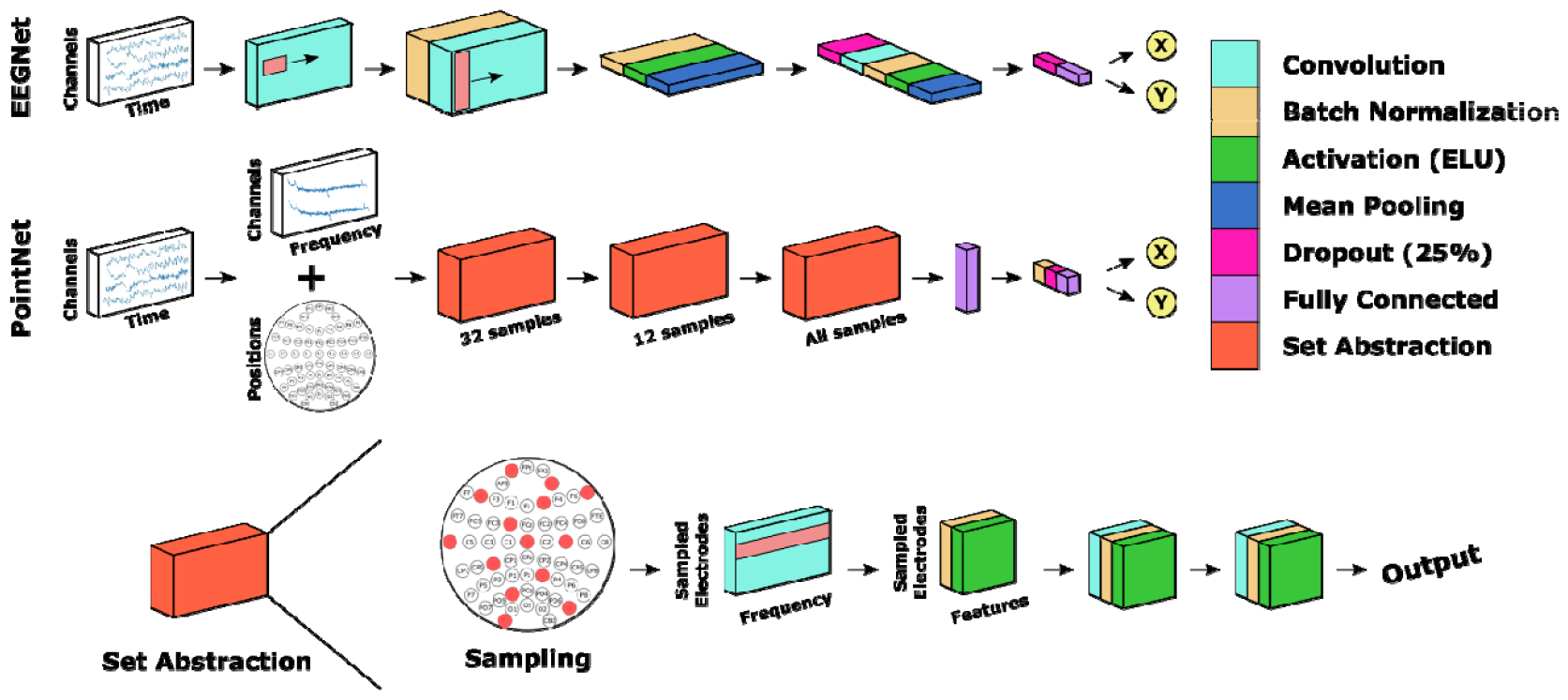
DL architectures. A) The implementation of EEGNet used in this study. B) Our proposed PointNet architecture, adapted for BCI decoding.

### H. Study Design

Two of the main goals of this study were to establish DL-based BCI decoders for the CP task, and to evaluate their performance in human subjects to evaluate the effects of additional training data. To accomplish this, we developed the study design shown in Figure 5. Subjects completed 8 sessions of CP BCI experiments. Each session consisted of 12 CP runs, where each run had 5 one-minute trials, with an additional run used to calculate the chance level (Figure 5B). This chance level was used to estimate a baseline performance when there is no control over the system (55). More information about the chance level can be found in the Supplementary Materials. Since the DL-based decoders require subject-specific training data before they can be used, the first session consisted entirely of runs using the traditional decoder. Each session after the first had four runs each of the traditional decoder, PointNet model, and EEGNet model (Figure 5C). The order of decoders was randomized for each session, but all four runs of each decoder were completed together as a block.

Subjects were required to have at least two sessions of BCI experience before starting this experiment. This requirement helped to ensure that subjects had some control over the BCI system, reduce the effects of training on performance in the first few sessions, and prepare subjects for the more complex CP task. In addition, the speed of the cursor and target was reduced for the first session of this experiment (Session ‘T’ in Figure 5B) to help introduce subjects to the CP task. The speed was then increased and kept consistent for the remaining 7 sessions (Sessions 1-7 in Figure 5B).

### J. Transfer Learning

We designed a second sub-study to investigate the potential benefits of using transfer learning to reduce the amount of training data needed for DL-based decoders. For this experiment, we chose to test only the EEGNet architecture to allow us to collect more data for each experimental condition instead of having to split the runs between EEGNet and PointNet. We designed the study as shown in Figure 5C, similar to the original study design with 12 CP runs and an additional chance run for each session. Here, we recruited only subjects with no prior BCI experience and ran two sessions of DT BCI as training before moving on to the CP tasks. All of the subjects recruited for this experiment were different from those who participated in the previous CP experiment. We also chose to reduce the cursor speed by 25% and the variance of the random accelerations applied to the target by 33% for this experiment after seeing that some subjects struggled with the faster speeds in the first part of this study.

For this follow-up experiment, we tested the original EEGNet DL-decoder paradigm (DL) along with two new DL-based paradigms: transfer learning (TL) and recalibration (CL). Details on the on our implementation of transfer learning can be found in the supplemental materials.

## Data and code availability

The EEG BCI data in 28 human participants will be made public via Figshare when the paper is accepted for publication. The implementations of both the EEGNet and PointNet architectures used in this study will be made public via GitHub when paper is accepted for publication.

## Acknowledgments

The authors thank Jeehyun Kim for assistance with data collection.

