## Supplementary Materials for "Continuous Tracking using Deep Learning-based Decoding for Non-invasive Brain-Computer Interface"

Of

**Supplementary Methods:**

A. **DL Training paradigm**

Training was performed using the Adam optimizer (45) for 15 epochs. At the end of training, the model weights from the epoch that produced the best validation performance were used as the final model. Loss was calculated as the Mean Squared Error (MSE) between the x and y coordinates of the prediction and label, with an additional term of the KL-divergence between a Gaussian distribution of zero mean and unit standard deviation (Equation 1). Here, *pred* represents the model predictions, *label* is the data labels, and *I* is the identity matrix. This KL-divergence term rewards the model for having an output distribution resembling a Gaussian centered at zero, encouraging it to produce evenly distributed outputs. This is similar to the normalization step in the traditional decoder and is important for online BCI trials, where favoring one direction can cause the cursor to become stuck on one edge of the screen and render the data useless for training future models.

$\text{Loss=MSE(pred, label) + KL(pred, N(0,I))}$	(1)

For training data, we used 75% of the available data for training and the other 25% was held out as a validation set for choosing the best model weights. We did not use a test dataset since each of the models was evaluated in online BCI experiments after training. To split the data, every fourth run was used as validation data, while the other three runs were used as training data (this corresponds to the first three runs for each decoder being used as training data and the last as validation, see the study design in Figure 3 for more information).

The EEG signal, cursor positions, and target positions were recorded from online experiments. To calculate the label for supervised learning, we cut the data into 1 second windows (the length of the input for the DL models). The cursor positions and target positions throughout this 1s window were then averaged to obtain a single cursor position and a single target position for each data window. We then subtracted the cursor position from the target position (both two-dimensional vectors) to obtain a single 2D vector that points from the average cursor position for the window to the average target position for the window. Finally, we normalized this vector to unit length. This 2D vector (one horizontal component and one vertical component) was then used as the label for training the DL models.

**EEGNet** **-** The details of the EEGNet architecture can be found in the original publication: (50). Our implementation of EEGNet is shown in Figure 6A in the main manuscript. Instead of applying a SoftMax to the output, our implementation of EEGNet performed regression and produced two numbers, one corresponding to the horizontal component of the cursor-to-target vector, and another for the vertical component.

A window of 1 second of EEG data is used as the input as a 62 channel by 250-time points array (1 second of data at 250 Hz). The data in this window is downsampled by a factor of 4 (from 1 kHz to 250 Hz) and common average re-referenced. This preprocessing is done in both online experiments and offline training. In the online experiment, new data blocks arrive every 40ms, so the input window consists of the most recent 1 second of data, updated every 40ms. In the offline setting, data windows are sliding windows 1 second long, sliding every 40ms to stay consistent with the online setting.

**PointNet -** Unlike the EEGNet implementation, for PointNet we used a spectral domain representation of the EEG signal for the input to the model. The same data windows described above for EEGNet were used, but were passed through a Fast Fourier Transform (FFT) to yield an estimation of the frequency spectrum of the signal for each electrode. The input to the PointNet model is the array of 62 frequency spectrums (one for each channel), and a 62 x 3 vector containing the 3D locations of each electrode relative to one another. The positions were a set of standard electrode positions for the Neuroscan Quik-Cap, and were not recorded experimentally. The same electrode positions were used for every subject.

B. **Chance level**

Chance level is a commonly used benchmark in BCI studies to show the performance level when subjects have no control over the system (53). To estimate the chance level for our implementation of the CP paradigm, we ran an additional run in each session of the study where the screen was turned off and subjects were instructed not to perform any motor imagery. Subjects were also asked to relax and limit their movements to reduce any motion artifacts in the data. Since the subjects could not see the cursor or target and should not have performed any motor imagery, the performance from these runs should be random and can be used to estimate a chance level for this task.

C. **TL Training paradigm**

To implement transfer learning, we first trained an EEGNet model using the data collected from all 15 subjects in the first part of this study. A similar train-validation split was performed where the first 3 runs from each decoder block were used as training data and the fourth was kept for validation. Using this method, all subjects and sessions were represented in both the training and validation sets. The weights from this trained model were saved and referred to as the *base model*. By pretraining the EEGNet model on BCI data from previous subjects, we hypothesized that the models would need less subject-specific data to perform well.

This base model was used as the transfer learning model (TL in Fig. 5D) for each subject in the first session, before any subject-specific training data was available. We also chose to use the traditional decoder in the first session to compare with the performance of the base model. After every session, a new TL model was trained by initializing the weights to the base model weights and training using all available subject-specific data. For example, to create the TL model for Subject 1 Session 4, we started with a copy of the base model and fine-tuned it using Subject 1’s data from Sessions 1-3. This allowed us to use the base model as a starting point and refine it using subject-specific data to personalize the models.

The table below summarizes the different deep learning model training paradigms:

| Model Name | Initial Weights | Training Data | When to update |
| --- | --- | --- | --- |
| DL | Untrained weights | All previous sessions | Before the session |
| TL (Transfer Learning) | Base model* | All previous sessions | Before the session |
| CL (reCalibration) | Trained TL model | TL runs from only the current session | Mid-session, during the ‘chance’ runs |

*The “base model” was an EEGNet model trained using all of the data from the 14 subjects in the first sub-study. This model was used as the starting point for transfer learning and was fine-tuned using subject-specific data to create the TL models.

### **D. Recalibration**

The second new decoding paradigm we evaluated was recalibrating the DL decoders. By recalibrating the DL-based decoders mid-session with data from the current BCI session, we hypothesized that we could reduce the effects of inter-session variance in the data and achieve higher performance in the CP task.

In this method, subjects first performed 4 runs with TL decoder as normal. We then used those 4 runs from the current session to update the TL decoder mid-session. To do this, we created a new EEGNet model and initialized the model weights as the weights from the current TL decoder. We then trained the new model for 5 epochs using only the 4 runs of TL data from the current session as training data. To make the most of the data from the current session, all of the data from these 4 TL runs was used as training data and no validation was performed. The model weights at the end of the fifth epoch were used as the final weights for the recalibrated model (CL).

To train the CL model mid-session, we ran the random chance runs directly following the TL runs each session. While the chance runs were ongoing, the data from the TL runs were uploaded to a remote server, used to train the CL model, and then the resulting model weights were downloaded to the local computer to be used for the CL runs. Training the CL models with this setup took a similar amount of time as the chance-level run, so there was minimal interruption in the sessions.

**E. Traditional Decoder**

The EEG signal for the traditional decoder was processed in 160ms windows shifted every 40ms. First, a small surface Laplacian filter was applied to channels C3 and C4 by subtracting the average signal from the four surrounding electrodes. The filtered signals were then processed in an autoregressive spectral estimation model to extract a 3 Hz band from 10.5 to 13.5 Hz, the upper portion of the alpha (or mu) band. The sum of this band in C3 and C4 was then used in a linear classifier to produce an output in the horizontal (C4-C3) or vertical (-C3-C4) direction. Lastly, the most recent 30 seconds of classifier outputs were saved to estimate the mean and standard deviation of the control signal and to normalize the decoder output to around zero mean and unit standard deviation for each axis independently. This normalized signal was then used as the output to control the cursor velocity.

**F. Statistical Analysis**

Cursor and target position data were recorded using BCI2000 every 40ms and processed offline using custom MATLAB scripts. Statistical analysis was performed using custom R scripts. For offline analysis, the data was downsampled by a factor of 4 to 250 Hz and common average re-referencing was applied. Performance in the CP task is primarily quantified using either the error between the cursor and target, or the correlation between their positions. Here, we report cursor-target distance using the Mean Squared Error (MSE) between the two positions, normalized by the screen size (diagonal distance of the square workspace). The Pearson correlation coefficients are used to report the linear relationship between cursor and target positions throughout the trial.

For each of the NMSE, correlation, and ADA performance metrics, statistical analyses were performed by fitting linear mixed effects models. For these models, the session number, decoder, and session-decoder interaction were considered fixed effects while subjects were classified as random effects. ANOVAs performed on each of the models for NMSE, correlation (x and y), and ADA showed significant effects for session, decoder, and session-decoder interactions. Estimated marginal means, calculated from the R library *emmeans*, were used to perform pairwise t-tests to compare decoders in specific sessions (example Figure 2B). Unless noted otherwise, the Holm method was used to adjust p-values for multiple comparisons. Numerical p-values can be found in Section H of the Supplementary Materials.

The ADA metric was primarily used to compare DL models offline to choose the best architectures and weights for online experiments. To investigate how similar the ADA metric compares to cursor-target NMSE, we calculated the Pearson correlation coefficient between the two metrics for all trials. Furthermore, a linear model was fitted to the data with the NMSE as the dependent variable and ADA as the independent. The resulting parameters and linear fit are shown as results in Figure 2I.

To study the lag between target and cursor movements and the effect on performance, we constructed histograms of trials whose best performance was achieved at specific time-lags. To create these plots, we shifted the cursor position vector relative to the target position vector, with the difference between vector positions called the *lag*. Since the cursor and target positions were sampled every 40ms, each shift of the cursor position vector represents a lag of 40ms. After each shift, the NMSE between the cursor and target positions was calculated and recorded for that specific lag. The lag that produced the best NMSE for each trial was recorded and the trials were binned based on their best lag to construct histograms. Finally, the average NMSE across trials was found for each decoder at each lag and used to determine which lag resulted in the best average performance for each decoder.

**G. Computational resources and training times**

All DL decoders were trained on a custom-built GPU server (Exxact Corporation). This server had the following specifications: 2 24-core AMD EPYC 7413 processors, 512 GB of RAM, 7 NVIDIA RTX A5000 GPUs with 24 GB VRAM. Data were copied to the server after each experiment and the DL models were trained remotely on this GPU server. The resulting model weights were then downloaded to be used for the next session. For mid-session training (recalibration models), data were uploaded to the server at the start of the 5-minute chance-level runs, models were trained during this time, and the resulting weights were downloaded as soon as they were available to be used in the same session.

BCI experiments were performed on one of two identical desktop workstations with the following specifications: Dell XPS 8930 with a 6-core Intel i7-8700, 64GB of RAM, and an NVIDIA GeForce GTX 1070 Ti.

To obtain approximate training times after all experiments were completed, we trained one model for each decoder using 7 sessions of data. A single A5000 GPU was used in each case. Using this setup, EEGNet took about 2 hours to train, while PointNet took about 40 minutes. EEGNet recalibration times took approximately 5-5.5 minutes, including the short pre-processing scripts that are required to cut data into appropriate windows and add the labels. The exact training time for the base model for transfer learning was not recorded, but this training took several days to complete.

Approximate inference times were estimated by recording the inference time for each update window during one run of an online experiment (about 7500 predictions, 25 predictions per second multiplied by 300 seconds of BCI control) and taking the mean. Using this method, EEGNet had a mean inference time of 4.57ms with a standard deviation of 1.19ms. PointNet had a mean time of 9.8ms and a standard deviation of 1.90ms. These inference times were calculated by running the CP task on the Dell XPS desktop computers that were used in the main study.

**H. Numerical p-values**

**Figure 2B**

| **Decoders** | **P-value** |
| --- | --- |
| ANOVA across Decoders | <0.0001 |
| Chance / AR | <0.0001 |
| Chance / EEGNet | <0.0001 |
| Chance / PointNet | <0.0001 |
| AR / EEGNet | 0.7019 |
| AR / PointNet | 0.0052 |
| EEGNet / PointNet | 0.1134 |

**Figure 2C**

| **Decoders** | **P-value** |
| --- | --- |
| ANOVA across Decoders | <0.0001 |
| Chance / AR | <0.0001 |
| Chance / EEGNet | <0.0001 |
| Chance / PointNet | <0.0001 |
| AR / EEGNet | <0.0001 |
| AR / PointNet | <0.0001 |
| EEGNet / Point | 0.2768 |

**Figure 2G**

| **Decoders** | **P-value** |
| --- | --- |
| ANOVA across Decoders | <0.0001 |
| Chance / AR | <0.0001 |
| Chance / EEGNet | <0.0001 |
| Chance / PointNet | <0.0001 |
| AR / EEGNet | 0.8328 |
| AR / PointNet | 0.4515 |
| EEGNet / Point | 0.9216 |

**Figure 2H**

| **Decoders** | **P-value** |
| --- | --- |
| ANOVA across Decoders | <0.0001 |
| Chance / AR | <0.0001 |
| Chance / EEGNet | <0.0001 |
| Chance / PointNet | <0.0001 |
| AR / EEGNet | <0.0001 |
| AR / PointNet | <0.0001 |
| EEGNet / Point | 0.6182 |

**Figure 4B**

| **Decoders** | **P-value** |
| --- | --- |
| ANOVA across Decoders | <0.0001 |
| Chance / AR | <0.0001 |
| Chance / Recalibration | <0.0001 |
| Chance / Transfer Learning | <0.0001 |
| AR / Transfer Learning | 0.6224 |
| AR / Recalibration | 0.8597 |
| Transfer Learning / Recalibration | 0.9753 |

**Figure 4C**

| **Decoders** | **P-value** |
| --- | --- |
| ANOVA across Decoders | <0.0001 |
| Chance / DL | <0.0001 |
| Chance / Recalibration | <0.0001 |
| Chance / Transfer Learning | <0.0001 |
| DL / Transfer Learning | 0.3723 |
| DL / Recalibration | 0.8331 |
| Transfer Learning / Recalibration | 0.0676 |

**Supplementary Figures:**

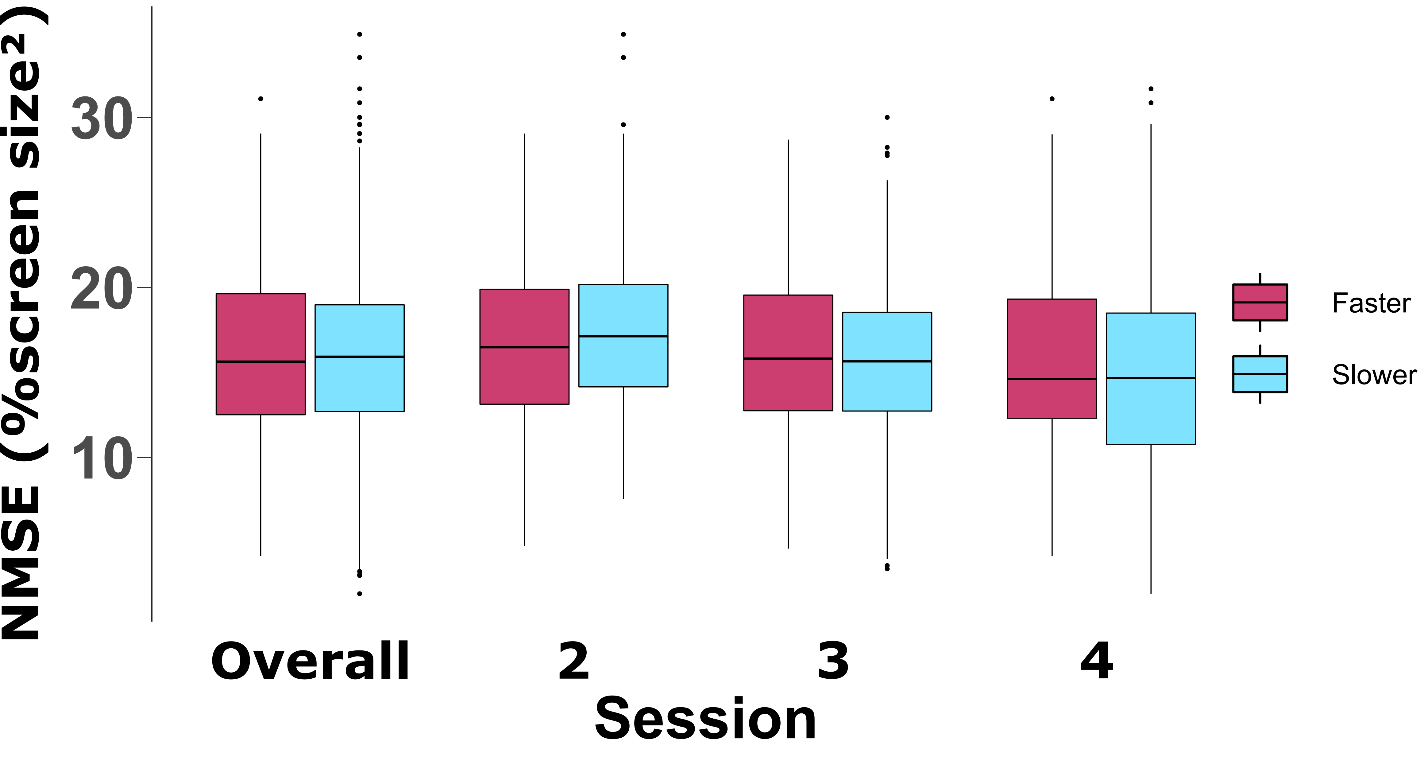

**Supplementary Figure S1.** Comparison of speeds from the first and second studies. Since EEGNet was the only model used in both experiments, only EEGNet (no transfer-learning) runs are included here. This corresponds to the ‘EG’ decoder in the first study and the ‘DL’ decoder in the second. A linear mixed-effects model was fit to the data with speed, session, and the interaction between speed and session as the fixed effects, and the subject as the random effect. Although the speed was reduced in the second study to help improve subjects’ performance, an ANOVA on the fitted model found neither speed nor the interaction between speed and session to have a significant effect on performance.

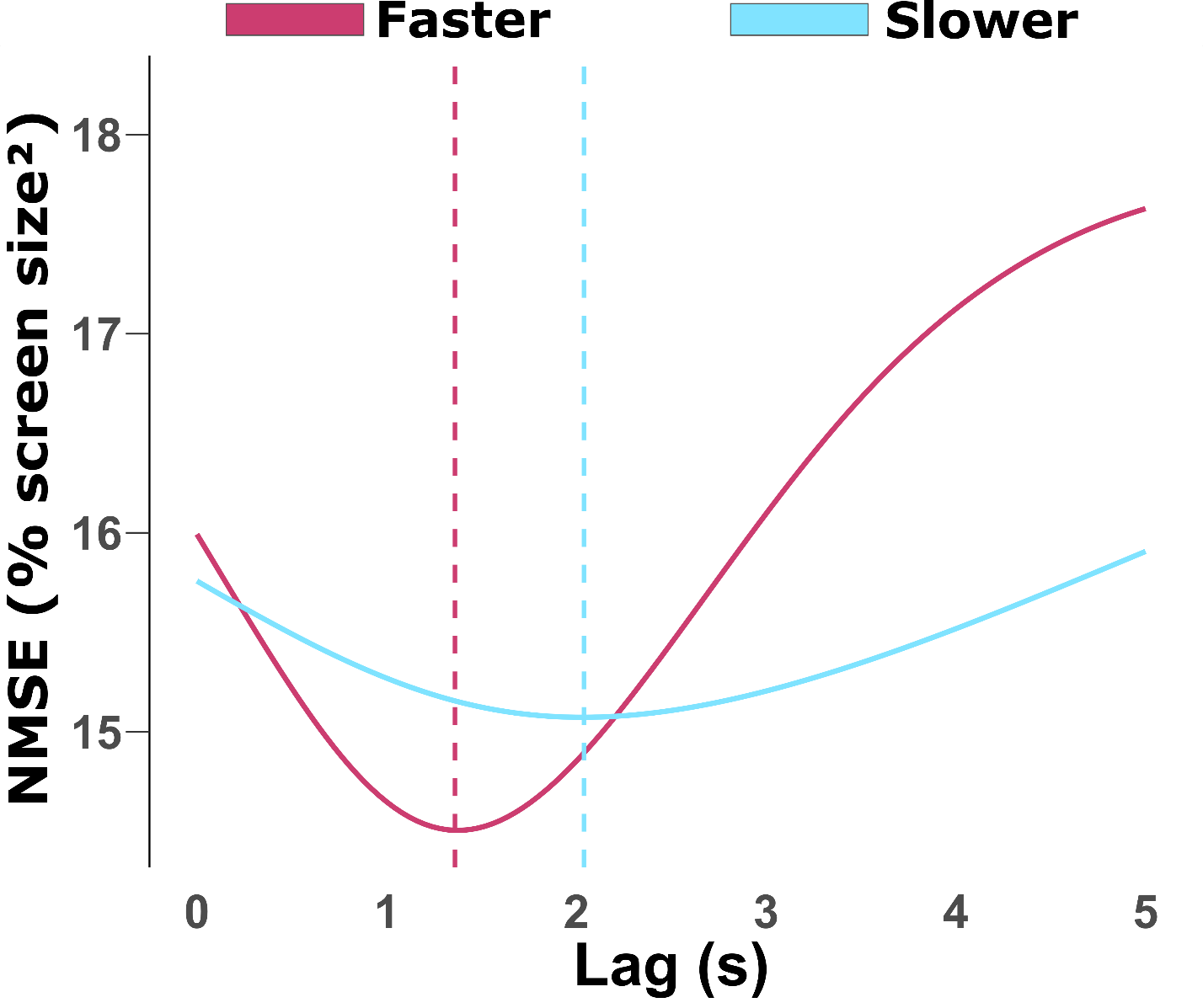

**Supplementary Figure S2.** Effects of lag on MSE performance for both the faster and slower speeds. Here, only the EEGNet runs were analyzed (‘EG’ in the first study and ‘DL’ in the second) since this was the only paradigm used in both sub-studies. Although the best lag for the slower speed occurs at a later time (2.04s), there is less of a change in performance than the faster speed (peak at 1.34s). Based on these results, the lag phenomenon appears to affect the faster speeds more but still decreases performance somewhat even when using slower speeds. One explanation for this result could be that the slower speed allows subjects more reaction time to change their motor imagery, but the slower speed also increases the time it takes for them to correct their course.
